# Strong population structure in Venezuelan populations of *Coccidioides posadasii*

**DOI:** 10.1101/719328

**Authors:** Marcus M. Teixeira, Primavera Alvarado, Chandler Roe, George R. Thompson, José Salvatori Patané, Jason W. Sahl, Paul Keim, John N. Galgiani, Ana Litvintseva, Daniel R. Matute, Bridget M. Barker

## Abstract

*Coccidioides posadasii* is a pathogenic fungus that causes coccidioidomycosis in many arid regions of the Americas. One of these regions is bordered by the Caribbean Sea, and the surrounding landscape may play an important role in the dispersion of *C. posadasii* across South America through southeastern Mexico, Honduras, Guatemala and Venezuela. Comparative phylogenomic analyses of *C. posadasii* reveal that clinical strains from Venezuela are genetically distinct from the North American populations found in Arizona (AZ), Texas, Mexico, and the rest of South America (TX/MX/SA). We find evidence for admixture between the Venezuela and the North American populations of *C. posadasii* in Central America. As expected, the proportion of Venezuelan alleles in the admixed population decreases as latitude (and distance from Venezuela) increases. Our results indicate that the population in Venezuela may have been subjected to a recent bottleneck, and shows strong population structure. This analysis provides insight into potential for *Coccidioides* spp. to invade new regions.

## INTRODUCTION

In spite of their human health impact, fungal pathogens are currently understudied (1, 2). To date approximately 150 species are responsible for disease in humans, with an estimated 1.5 million deaths each year (3, 4). The most affected are immunocompromised patients (e.g. HIV), but several of these diseases can also affect otherwise healthy patients (5, 6). Systematic surveys of genetic diversity of fungal pathogens have revealed extensive variability in the strength of virulence among genotypes of isolates from the same species (e.g., (7)). Other traits like antifungal resistance, and ability to survive different environmental conditions also show extensive variation (8–10). Understanding the magnitude and sources of variation among these traits is a crucial aspect of understanding why some pathogens are more effective at spreading and causing disease than others.

An important aspect of variation is not only the total magnitude within a species but also how that variation is apportioned across populations (11, 12). Population structure within species and barriers to gene flow can maintain genetic variation within and between species boundaries, respectively. Studying the origin and maintenance of genetic variation is important because it is likely to dictate the pathogen’s ability to cause disease and respond to management strategies (13). The paucity of studies that have focused efforts to understand the magnitude of this variation is particularly acute for fungal pathogens that occur in the tropics and subtropics. The tropics are generally more diverse than temperate areas in terms of numbers of species (14–16), thus the genetic diversity of fungi, and in particular of fungi with the ability to cause disease in humans, might be larger in the tropics as well.

Coccidioidomycosis or “Valley Fever” is an example of one of these primary fungal diseases, and is caused by the saprobic species *Coccidioides immitis* and *C. posadasii* (17). The disease range overlaps arid regions throughout the American continent, but California and Arizona encompass the vast majority of reported cases of the disease (18). Due to the availability of many clinical isolates, most sampling has occurred in these two US states, even though the geographic range extends into Latin America. Coccidioidomycosis has been reported less frequently in arid and semiarid environments of Central America (Guatemala and Honduras) and South America (Brazil, Argentina, Paraguay, Colombia and Venezuela) (19, 20). It is unclear if isolates of *Coccidioides* from the southwestern United States show a pattern of higher infectivity and/or virulence, or if it reflects a lower prevalence of the organism in other endemic regions in the Americas coupled with lower population densities in most arid regions. Alternatively, the pattern could be explained by lower rates of fungal disease awareness, testing, and reporting from tropical regions.

The two etiological agents of coccidioidomycosis, *C. immitis* and *C. posadasii*, are two sibling species that diverged about 5 million years ago (21). The species are thought to be phenotypically similar but some key differences have been reported. Besides showing a highly differentiated genome, *C. posadasii* grows more slowly than *C. immitis* at high concentrations of salt (22). Additionally, spherules of *C. posadasii* appear to develop asynchronously as compared to *C. immitis* (23). There are also broad differences in the geographic distribution of the two species. While *C. immitis* is restricted to North America, *C. posadasii* has a broader distribution, which ranges from Arizona to Argentina. Consistent with this distribution, coccidioidomycosis in South and Central American is primarily caused by *C. posadasii* but reports of *C. immitis* exist from Argentina and Colombia (24–26). Both species of *Coccidioides* show strong signatures of population structure. In the case of *C. immitis*, genomic analyses have revealed the existence of two subdivided populations: Washington state (at the northernmost point of the known range of the species distribution) and the rest of the range (27). In the case of *C. posadasii* at least two distinct populations have been proposed: Arizona (AZ) and Texas/Mexico/South America (TX/MX/SA) (21, 28).

Studying population structure of *C. posadasii* is critical to understanding the spread of the pathogen across the Americas. Based on microsatellite analyses, the introduction of *C. posadasii* into South America has been dated between 9,000 to 140,000 years ago, and was proposed to follow the Amerindian intercontinental migration (29). However, the most recent common ancestor for *C. posadasii* populations, inferred by whole genome SNP analysis, revealed that *C. posadasii* lineages that gave rise to TX/MX/SA and AZ emerged 700kya, and *C. posadasii* Guatemala about 200kya (21). These two findings are not consistent with a serial bottleneck giving rise to the current South American populations, but are consistent with ancestral population structure previous to the spread of *C. posadasii* to South America and the migration of a few lineages to the south. However, before any of these hypotheses can be tested, an assessment of the genetic characteristics of South American isolates is sorely needed.

One of these most characteristic xeric ecosystems in South America is the Paraguaná xeric shrubland (30), which is characterized by arid and dry climates, low altitude, xerophytic vegetation, and sandy soils with high salt. This environment may favor *Coccidioides* spp. growth and development (31). Notably, skin test surveys using coccidioidin in Latin American communities revealed positive testing in 44% and 46% in the Chaco region of Paraguay and the Lara state of Venezuela, respectively (26, 32). Additionally, environmental molecular detection of *Coccidioides* spp. in Venezuela suggests a high prevalence of *C. posadasii* in the soil (33). The actual occurrence of coccidioidomycosis remains unclear; as fewer than 1,000 total coccidioidomycosis cases have been reported over the last century in South and Central America (19, 26). However, environmental sampling and serological inquires suggest that the importance and prevalence of coccidioidomycosis in South America may be vastly underappreciated, and little is known about the genetic characteristics and genealogical relationships of this population.

In this report, we aim to bridge this gap. We sequenced 10 *C. posadasii* isolates from Venezuela and studied their relationship to other isolates. These isolates form a monophyletic group with little diversity, which is differentiated from other *C. posadasii* populations. Notably, we find that Central American populations of *C. posadasii* are the result of admixture between North America and Venezuela. These results reveal the importance of characterizing tropical populations of *Coccidioides* as they harbor distinct genetic variants, and likely phenotypic differences, among populations. These populations can act as donors of variation, and contribute to the evolution of other populations in subtropical and temperate regions.

## MATERIALS AND METHODS

### Fungal strains, DNA extraction and DNA sequencing

#### *Coccidioides* isolates

*Coccidioides* isolates were retrieved from human clinical specimens collected at the at the Servicio Autónomo Instituto de Biomedicina Dr. Jacinto Convit, Caracas, Venezuela; the Valley Fever Center for Excellence, Tucson, Arizona; or at the UC Davis Center for Valley Fever, UC Davis Health, Davis, California. Clinical specimens were initially seeded onto Mycosel agar (BD Biosciences), or Sabouraud agar with 100 ug/mL chloramphenicol and cycloheximide, and fungal isolates were subjected to serial passage onto 2X-GYE media (1% w/v Difco yeast extract and 2% w/v glucose) for fungal colony stabilization and to detect any potential contamination. *Coccidioides* arthroconidia were harvested and preserved in 25% glycerol, 0.5% glucose, and 0.25% yeast extract storage media, and finally stored at −80°C.

### DNA extraction

To culture isolates and extract DNA, we grew *Coccidioides* (isolates listed in Supplementary Table 1) in BSL-3 conditions either at the Pathogen and Microbiome Institute, University of Northern Arizona, Flagstaff, Arizona, USA or at the Servicio Autónomo Instituto de Biomedicina Dr. Jacinto Convit, Caracas, Venezuela. We seeded all fungal cultures from glycerol stocks, previously kept at −80°C, onto 2X-GYE media for mycelia propagation at 28°C for 14 days. We harvested ~500mg of mycelia using a cell scraper (VWR, Radnor, PA) and this material was used as input for total DNA extraction using the UltraClean Microbial DNA Isolation Kit (Qiagen) according to the manufacturer’s protocol. We then confirmed the sterility of all DNA before removing the DNA preparations from the BSL3 facility for further sequencing by plating 5% of the eluted material onto 2X-GYE media and growth was evaluated after 72 hours at 28°C. Once removed from the BSL3, we estimated DNA purity and concentration using spectrophotometry on the NanoDrop^®^ ND-1000 system (Thermo Fisher Scientific).

### Library preparation and sequencing

We prepared sequencing libraries for the ten *de novo* sequenced isolates using the Kapa Biosystems kit (Kapa Biosystems, Woburn, MA) and ~1μg of DNA of each isolate according to manufacturer’s protocols. We then multiplexed individual libraries using 8-bp indexes and quantified using quantitative PCR (qPCR) in a 7900HT system (Life Technologies Corporation, Carlsbad, CA) using the Kapa library quantification kit (Kapa Biosystems, Woburn, MA). We pooled libraries that were quantified using qPCR as above, and sequenced on an Illumina HiSeq 2500 instrument (Illumina, San Diego, CA) at the Translational Genomics Research Institute (TGen) aiming for a coverage of 100X per isolate. Resulting paired end reads had a length of 125bp.

### Public data

We obtained 72 publicly available sequence reads for *C. posadasii* (51) and *C. immitis* (21) isolates from the Sequence Read Archive (SRA). All accession numbers are listed Supplementary Table 1.

### Read mapping and variant calling

First, we removed Illumina adaptors using Trimmomatic v 0.36 from the reads obtained for the 10 sequenced isolates (34). To improve *de novo* references, we used the Unmanned Genome Assembly Pipeline (UGAP – https://github.com/jasonsahl/UGAP) that uses the genome assembly algorithm SPAdes v 3.10.1 (35) as well the Pilon toolkit v 1.22 (36). We obtained publicly available *Coccidioides* raw reads from the Sequence Read Archive (SRA) deposited at the accession number SRR3468064 (C. *posadasii* Nuevo Leon-1) and SRR1292225 (C. *immitis* strain 202). We used Burrows-Wheeler Aligner (BWA) v 0.7.7 (37) to align reads to each of the assembled references: *C. posadasii* strain Nuevo Leon-1 (27) or *C. immitis* strain 202 (38). We removed mismatching intervals with the RealignerTargetCreator and IndelRealigner tools available in the GATK toolkit v 3.3-0 (39, 40). To call SNPs, we used UnifiedGenotyper setting the parameter “het” to 0.01. Finally, we filtered the .vcf files using the following parameters: QD = 2.0 || FS_filter = 60.0 || MQ_filter = 30.0 || MQ_Rank_Sum_filter = −12.5 || Read_Pos_Rank_Sum_filter = −8. SNP’s with less than 10X coverage or with less than 90% variant allele calls or that were identified by NUCmer (41) as being within duplicated regions in the reference were removed from the final dataset. In total, our dataset was composed of 78 *Coccidioides* genomes.

### Phylogenetic tree

To study the genealogical relationships among *C. posadasii* isolates, we built a Maximum Likelihood (ML) phylogenetic tree using genome-wide SNP data. We used concatenated genome-wide SNPs as *Coccidioides* as we expect low levels of genealogical discordance (42, 43). We first obtained whole supercontig sequences for each individual from the VCF file using the FastaAlternateReferenceMaker tool in GATK. Next, we used IQ-TREE to infer the most likely tree with maximum likelihood (44); we used the -m TEST option (jModelTest (45)) for automatic molecular evolution model selection. We calculated the support of each branch on the resulting topology, using 1,000 ultrafast bootstraps coupled with a Shimodaira–Hasegawa-like approximate likelihood ratio test (SH-aLRT) (46, 47). Tree topologies were visualized using FigTree v1.4.2 – http://tree.bio.ed.ac.uk/software/figtree/.

### Mating type

We determined whether the genomes from the *C. posadasii* population from Venezuela have evidence of functional mating type loci based on sequence similarity to known mating type sequence. The Onygenalean mating-type locus contains one of two forms of unrelated sequences (known as the idiomorph) in a syntenic but unique segment that gives sexual identify to fungal cells. To identify the mating type of the Venezuelan *C. posadasii* isolates we followed the same strategy previously used to identify the *MAT1-1* or the *MAT1-2* loci in this pathogen (27, 48). Briefly, we used the full *MAT1-1* (containing the alpha-box *MAT1-1-1* gene – EF472259.1) and *MAT1-2* (containing the HMG-box *MAT1-2-1* gene EF472258.1) loci as reference sequences to query the Venezuelan *C. posadasii* read files. All isolates examined to date carry only one of these loci, and are classified as either a *MAT1-1* or *MAT1-2* genotype.

### Approximate time of divergence

We used Beast v2.5.2 (49) to infer molecular dating estimates. The analysis was performed in two steps. In the first step, we run the program twice including all the taxa, but assuming two different time priors following a normal distribution truncated at zero (with means at 5.1 Mya and at 12.8 Mya respectively, each with a standard deviation = 10%). We assumed a birth-death tree prior. In the second step, we assumed a coalescent model (skyline; (50)) including the Caribbean population only (i.e., excluding the reference *C. immitis)* with root prior age extrapolated from the age ranges rather than 95% highest posterior densities (HPD) for this same node in runs of the first step, under a uniform distribution. We assumed the prior root age to be within the range given by the minima and maxima of the ranges considering the two different runs of the first step, i.e., [min(minima); max(maxima)]. After the second step was completed, we generated a skyline plot. For each scenario, we ran two MCMC runs until convergence was detected and effective sample sizes were > 200, as revealed in Tracer v1.7 (51). We then summarized the two runs for each case using treeannotator (within the Beast v2.5.2 distribution) after discarding the burnin regions, summarizing ranges and 95% HPD of divergence times.

### Population structure

We studied the population structure of *C. posadasii* using two different methods. First, we used Principal Components Analysis (PCA) which provides a graphical representation of the partition of genetic variance in a population sample. To do so, we restricted our dataset to biallelic sites and used the R package *‘adegenet’* (52). We used the function fasta2genlight to extract biallelic SNPs from VCF files (see above) and the function glPca to compute the first two principal components (PCs). We also report the percentage of genetic variance explained by each PC. Next, we inferred the most likely population clustering within *C. posadasii* using fastSTRUCTURE v1.0 (53). This tool calculates the most likely number of populations (K) and the probability that each strain belongs to each population. SNPs were assumed to be unlinked under the admixture model, and the ancestry of each individual and correlated allele frequencies were simulated for a range 2 to 8 populations. To infer the most likely true number of populations, we found the K with the lowest likelihood using the script chooseK.py (53), which parses the log files of each inferred value of K and infers the best fitting model by maximizing the marginal likelihood.

### Genetic distances

One of the proxies of cryptic speciation is that isolated species show considerably larger genetic differentiation than the magnitude of intrapopulation variation (43). We studied whether *C. posadasii* showed the signature of cryptic speciation. We measured the genetic differentiation between the nine groups defined by the above phylogenetic analysis. To calculate the between group distance, we used Dxy, the mean number of genetic differences between two genomes from different clusters. We compared the mean differentiation between two groups to the magnitude of genetic diversity within groups. To calculate the within group diversity, we used π, the mean number of genetic differences between two genomes from the same cluster. Genome-wide SNP data was loaded into MEGA7 (54) and the genetic groups were determined according to the groups revealed in the phylogenetic tree as follows: *C. immitis, C. posadasii* Venezuela, *C. posadasii* Guatemala, *C. posadasii* TX/MX/SA, *C. posadasii* AZ1, *C. posadasii* Phoenix, *C. posadasii* Tucson and *C. posadasii* Tucson24/3490/GT120/Sonora. For pairwise comparisons we use asymptotic 2-sample permutation tests using the function *perm.test* (R library ‘exactRankTests’). Since there were two values of π for each value of Dxy, we compared Dxy values to the highest value of π in a pair of populations. We adjusted the significance threshold to account for multiple comparisons to P = 0. 00138 (0.05/36).

### Admixture

Previous studies have found that species of *Coccidioides* exchange alleles (55, 56). Our scope was different; we assessed whether populations within *C. posadasii* exchange genes. We estimated the proportion of admixture for each isolate using *ADMIXTURE* (57). We inferred the number of populations within *C. posadasii* by testing which K scenario had the lowest marginal likelihood, in a similar manner to that described for fastSTRUCTURE (see above). In the case of the Caribbean population, we regressed the proportion of admixture to the distance from the putative donor population, Venezuela.

Next, we used f4 statistics to assess the proportion of admixture. We restricted this analysis to the Guatemala population. We used admixtools (Patterson et al. 2012) implemented in Treemix (58) and the option -k 1000 was used to group 1,000 SNPs to account for linkage disequilibrium. Assuming a given phylogeny, the f4 ratio allows estimating the two mixing proportions during an admixture event even without access to the precise populations that gave rise to the admixed lineage for the two ancestral populations. Since we have inferred the phylogenetic relationships among *Coccidioides* populations, we calculated the proportion Venezuela ancestry in Guatemala following the equation:

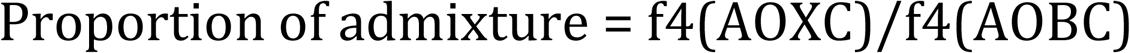

where X corresponds to the potentially admixed population (i.e., Guatemala). O, B A, and C are four populations that are known to branch at four distinct positions along the phylogeny. Outgroup refers to the outgroup; in this case, we used *C. immitis*. A and C are the donors (or close relative to the donors) of the admixed lineage. In this case, we used A = Phoenix, and C = Venezuela. B is a population that does not harbor introgression; we used B=Tucson. We found no difference in the estimations by swapping Tucson and Phoenix (data not shown).

Finally, we tested whether the contribution of Venezuela to the admixed population between Venezuela and Arizona followed the expectation that as distance from Venezuela increased, the contribution of the Venezuela population would decrease. Our hypothesis was that individuals from Central America were admixed and the product of gene exchange between the North American and South American *C. posadasii* populations. We used the proportion of admixture inferred from ADMIXTURE when K=3 in *C. posadasii* (the best fitting scenario, see results). Next, we calculated the distance between the collection site of a given sample (defined to the level of country or state) and Caracas using a haversine formula (59). Next, we calculated the distance between Caracas and the closest major city to where the isolate was collected using the great-circle distance between two points (i.e., the shortest distance over the Earth’s surface). We used the approximate coordinates of seven locations to calculate the waypoints distance from Caracas (10.4806° N, 66.9036° W). We used the following sites and coordinates: Guatemala (15.7835° N, 90.2308° W), Coahuila (27.0587° N, 101.7068° W), Arizona (34.0489° N, 111.0937° W), Florida (27.6648° N, 81.5158° W), Texas (31.9686° N, 99.9018° W), Michoacan (19.5665° N, 101.7068° W), and Sonora (29.2972° N, 110.3309° W). We used a one-tailed Spearman correlation test using the R package ‘stats’ (function ‘cor.test’).

## RESULTS

### Data availability and SNPs

All the raw reads for the 10 isolates we sequenced in this manuscript were deposited at SRA under BioProject number PRJNA438145 with sample accessions numbers SRR6830879-SRR683088 (Table S1). Reads from the 51 *C. posadasii* previously sequenced genomes were aligned to the reference *C. posadasii* strain Nuevo Leon-1. This alignment had 261,105 SNPs. When we included *C. immitis* 202 strain as the outgroup, the alignment had 464,281 SNPs. The difference between in the number of polymorphic sites between these two alignments reflects the genetic differentiation between *C. immitis* and *C. posadasii*.

### Phylogenetic tree and mating type analysis

First, we found that the model GTR+F+ASC+R9 was the best fit for the *C. posadasii* strain Nuevo Leon-1 alignment while TVM+ASC+G4 model was the best fit for the alignment for the *C. posadasii* and *C. immitis* 202 alignment. We then used these alignments and modes of molecular evolution to generate a maximum likelihood phylogenetic tree. We rooted the tree with *C. immitis*, the closest relative of *C. posadasii*.

The resulting maximum likelihood tree revealed that *C. posadasii* encompasses two major groups, one formed exclusively by Arizona isolates and one more heterogeneous that includes all the isolates from South America, central America, Texas, and a few isolates from AZ (Figure 1). The branches shown in Figure 1 have bootstrap and SH-aLRT support of over 90%. As expected the longest branch in the tree (i.e., genetic distance) is the branch that separates *C. posadasii* and *C. immitis*, the two species of *Coccidioides*. Consistent with previous studies (27, 56, 60), we estimate that the differentiation between these two species occurred 4.8 MYA (95% HPD – 3.8-5.8 MYA) years ago.

**FIGURE 1.**
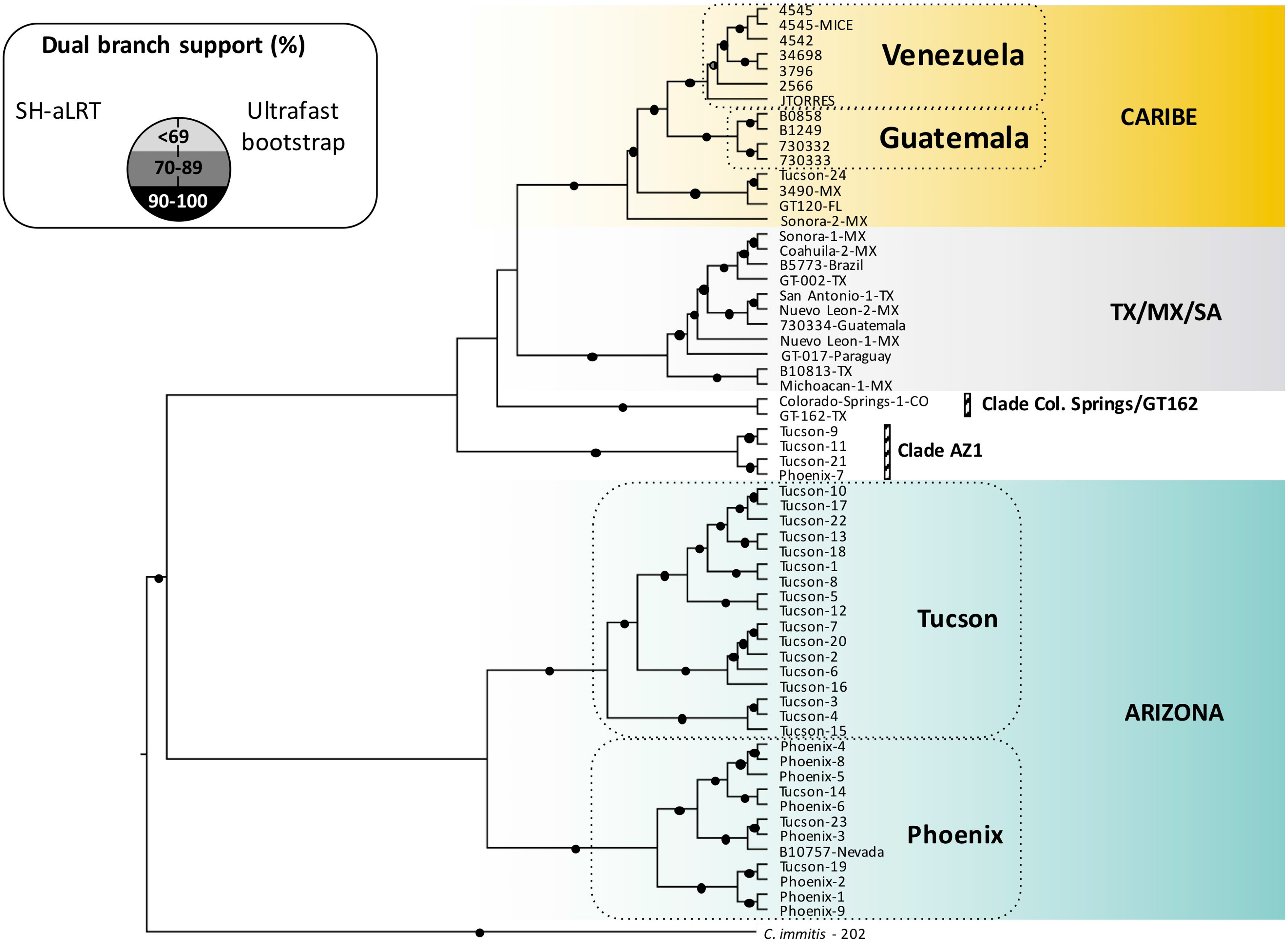
Maximum likelihood (ML) phylogenomic analyses among *C. posadasii* complex. ML tree was rooted with the *C. immitis* strain 202 and clades are displayed proportionally to the branch length of the clades since the majority of SNPs are derived from the overall *C. posadasii/C. immitis* divergence. Dual branch support was evaluated using 1,000 ultrafast bootstraps coupled with a Shimodaira-Hasegawa-like approximate likelihood ratio test and are displayed next to the clades.

The Arizona (AZ) clade contains two main groups, one composed of isolates from Tucson/Pima County, and one mostly (but not exclusively) composed by isolates from Phoenix/Maricopa County (Figure 1). These localities are both in the state of Arizona, USA. This level of micro geographic structure is puzzling because of the geographical proximity between the two sites (160 km), and because the majority of these isolates were collected from human patients. One would expect people to move freely between these two localities. This pattern might suggest that even though people can move between these two cities, there is a high level of predictability of where infections were contracted (i.e. the patient’s main residence).

The second clade encompasses a more diverse geographical sample that includes isolates from North, Central and South America (Figure 1). This group is composed of four subgroups. The first one is a clade formed by isolates mostly collected in localities surrounding the Caribbean Sea, including isolates from Guatemala and Florida. This is one of the longest branches within *C. posadasii*. Venezuela forms a monophyletic group which is sister to this Caribbean region clade. In effect, the Caribbean group is paraphyletic when Venezuela is not included in the topology (data not shown). The second group of the non-Arizonan isolates of *C. posadasii*, encompassing isolates from South America, appear nested within isolates from Texas and Mexico (henceforth referred to as TX/MX/SA – Figure 1). The existence of this group, which harbors South America and North American isolates, is consistent with (but does not uniquely support) the hypothesis that a genetically diverse group from North America underwent a population bottleneck while expanding south, and gave rise to genetically depauperate populations in South America (61). A third group is formed by 4 isolates from Arizona (AZ-Clade1 in Figure 1). Even though these clinical isolates were collected in the same localities as the isolates from the AZ population, they are not associated with this population. This suggests that these infections might have been acquired in a location different from southern Arizona (Tucson or Phoenix), or that additional phylogenetic clades remain to be defined with additional sampling. Finally, we find a small phylogenetic group composed of one isolate from Texas and one from Colorado (Figure 1). A more systematic sampling is needed to fully understand the genealogical and geographic relationships among *C. posadasii* groups, but the results from this genome-wide phylogenetic tree suggest there is differentiation among *C. posadasii* populations based primarily on geographic origin.

### Population structure

We next explored the partitioning of genetic diversity within *C. posadasii* using population genetics approaches. First, we assessed how genetic diversity was apportioned among isolates with a principle components analysis (PCA). We performed the analysis in two different ways. First, we included *C. immitis* and *C. posadasii* in the sample (Figure 2A). As expected, PC1 (75.34% of the variance) separates the two *Coccidioides* species. Consistent with previous findings, we see no intermediate isolates suggesting that even though admixture between the two species of *Coccidioides* has occurred (28, 62), the amount of genetic exchange between species is small enough to not affect species delineations. Notably, PC2 (3.03% of the variance) differentiates between the Arizona + TX/MX/SA and Venezuela populations with the Caribbean (e.g., Guatemala, Florida) group appearing as intermediaries between these two populations. PC2 thus corresponds to the population variation within *C. posadasii*. Interestingly with this analysis, the Guatemala population does not appear as a separate lineage but instead appears to be an intermediate between the Arizona and Venezuela populations.

**FIGURE 2.**
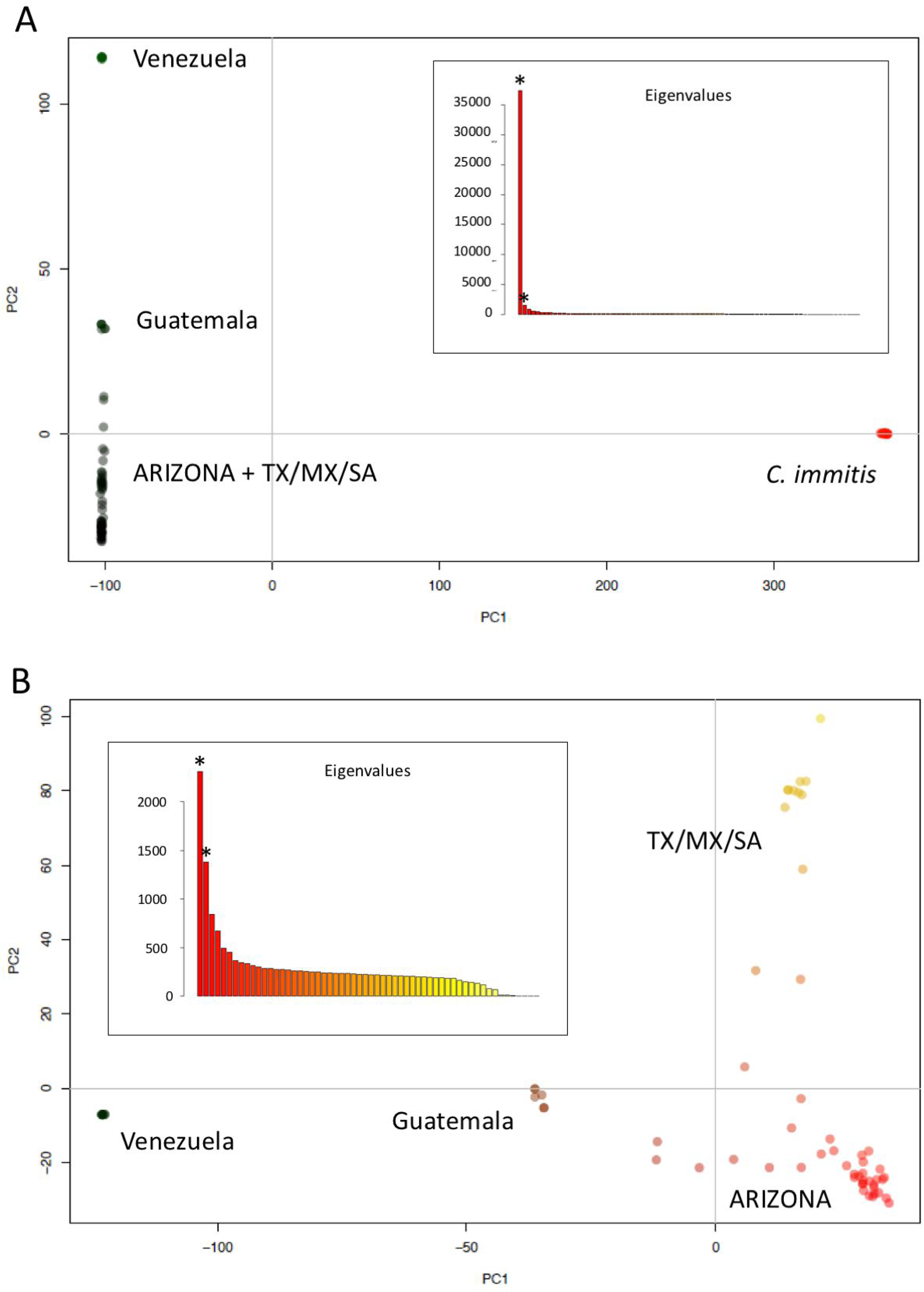
Principal component analysis (PCA) reveals population structure of *C. posadasii*. A) PCA including *C. immitis* and *C. posadasii*. PC1 separates between species of *Coccidioides*, while PC2 suggests the existence of cryptic populations within *C. posadasii*. B) PCA including only isolates from *C. posadasii*. PC1 coordinates separates isolates from Venezuela and isolates from the rest of the geographic range. PC2 discriminates the TX/MX/SA and ARIZONA populations. The insets on each panel show the eigenvalues that were used to plot the two principal coordinates (*represents the first 2 PCs).

The PCA that only included *C. posadasii* showed similar patterns but added more resolution to the differentiation within *C. posadasii* (Figure 2B). PC1 (14.26% of the variance) separates Venezuela and the remainder of the *C. posadasii* populations. PC2 (8.52% of the variance) separates the TX/MX/SA population from the rest of the *C. posadasii* Arizona populations. We find isolates that show genetic variation patterns that are intermediate between populations. The isolates from Guatemala and the isolates GT-120 (Miami, Florida), Tucson 7, and Tucson 2, appear as intermediaries between the AZ and Venezuela clusters (Figure 2B, PC1). This result is consistent with the PCA that included both species of *Coccidioides* in the previous analysis. Three isolates (Sonora 2, Michoacán 1, B01813-TX) from Mexico and Texas appear as intermediates between Arizona and TX/MX/SA (Figure 2B, PC2). These results are suggestive of population differentiation among populations between different geographical regions of *C. posadasii* and a degree of genetic exchange between these populations.

Second, we used fastSTRUCTURE with the admixture mode to infer the most likely number of groups in *C. posadasii*. When we include *C. immitis*, the method infers three clusters: *C. immitis, C. posadasii* from Arizona+TX/MX/SA, and *C. posadasii* from Venezuela (Figure 3). This result is qualitatively similar to the result from the PCA using the same dataset. When the analysis is run without *C. immitis*, we find a similar result. The genomic data reveal the existence of a Venezuela group, an Arizona group, and a third group formed by the TX/MX/SA isolates. The detection of the latter cluster is the main difference from the analysis run with *C. immitis*. These two results are consistent with the results the two PCAs (Figure 2) and suggest the existence of strong population stratification within *C. posadasii*. Notably, fastSTRUCTURE infers that the isolates from the Caribbean group, which appear as closely related to Venezuela in the phylogenetic tree, and as intermediates between Arizona and Venezuela in the PCA, are not as clearly assigned to a cluster. These show a high probability of being associated with the Venezuela group (P ≥ 0.85), but there is also a non-trivial probability they are associated with the Arizona group (P ≈ 0.15). The results are also largely consistent but not identical to previous attempts to determine the partition of genetic variation within *C. posadasii* (27); namely we find that Caribbean group is not an isolated population but instead is a population that cannot be assigned to either Venezuela or North America. The reasons for this conflict are explored below.

**FIGURE 3.**
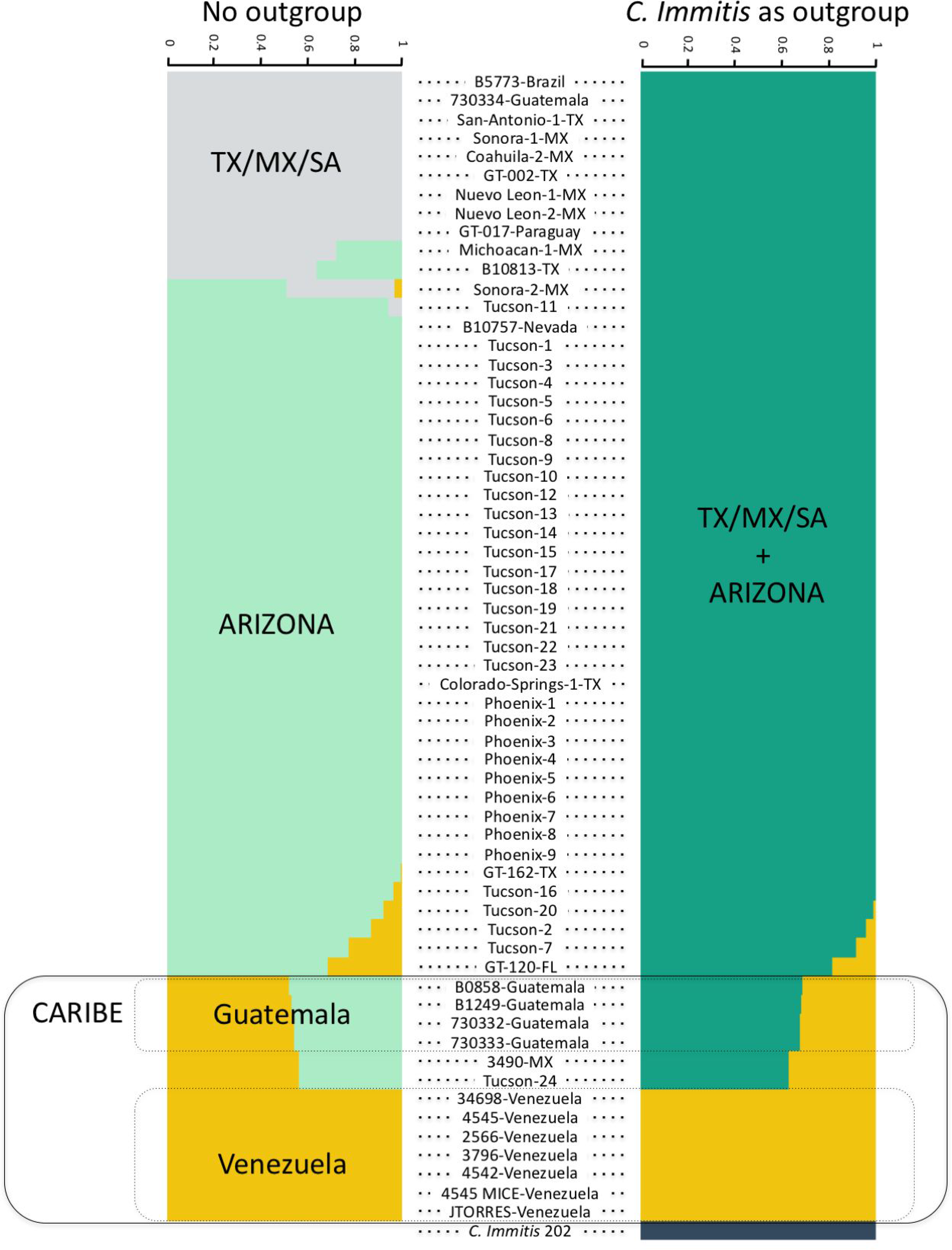
Strong population structure within *C. posadasii*. Population structure plots based on Bayesian posterior probabilities implemented in fastSTRUCTURE. We did the analysis in two ways, first using *C. immitis* as an outgroup for *C. posadasii* (A) or using only *C. posadasii* strains (B). Each row represents an individual. The height and colors of percentage of each population represent the probability of belonging to a given cluster.

### Genetic distances

Next, we studied whether there was evidence of cryptic speciation within *C. posadasii*. We measured the mean genetic distance between individual genomes in all pairwise comparisons to get a proxy of within population diversity and potential between population differentiation. Consistent with previous reports, *C. immitis* and *C. posadasii* are highly differentiated; the magnitude of interspecific differentiation is over 10 times larger than within species variation. The genetic distance value among populations is around 6% in all pairwise comparisons. Within each population, most π values are close to 5% (i.e. π_Tucson_ = 5.7%; π_Phoenix_ = 5.3%; π_TX/MX/SA_ = 4.5%). This value is in line with the levels of intraspecific variation described for other fungal species (63). However, the Caribbean group shows π values that are much lower. Heterozygosity in Guatemala is three times lower than in North American groups (π_Guatemala_ = 1.5%). A more extreme example is that of Venezuela, where heterozygosity is ~50 times lower than that of other *C. posadasii* populations (π_Venezuela_ =0.09%). We next compared these values to determine whether interpopulation pairs show a higher genetic distance than intrapopulation pairs. We find that this is the case in most comparisons (Table S2 showing results from asymptotic 2-sample Permutation Tests). These differences are consistent across groups and the magnitude of differentiation among populations is slightly higher than that within populations but does not clearly indicate the existence of cryptic species within the Caribbean group.

### Mating type

One possibility that might explain the low heterozygosity in the Venezuela population might be clonality due to the absence of one of the mating types, thus reducing recombination rates. All 7 isolates from the Venezuelan phylogenetic group harbor the *MAT1-2* idiomorph in their haploid genomes, while all other *C. posadasii* populations have balanced *MAT1-1* and *MAT1-2* idiomorphs (27). This prevalence of a single mating type is similar to observations for the clonal *C. immitis* Washington population (27), which suggests that population bottlenecks associated with range expansion might result in a single mating type and a clonal population structure.

### Admixture

PCA and fastSTRUCTURE results suggest that some isolates have mixed genetic ancestry between divergent *C. posadasii* populations. We studied the geographical partition of shared ancestry in the Caribbean group within *C. posadasii*. We used ADMIXTURE to infer the contribution from the minor population (i.e., Venezuela) as a proxy the proportion of admixture in each of the Caribbean group isolates. The results from ADMIXTURE show a similar clustering to the one revealed from fastSTRUCTURE and also suggest the presence of three populations within *C. posadasii* (Figure S1). The contributions of each of the three resulting populations to the admixed individuals of *C. posadasii* are shown in Figure 4. Next we studied the broad distribution of Venezuela ancestry across geography. Using the inferred proportion of Venezuela ancestry, we tested whether there was a relationship between the proportion of Venezuela contribution and the distance to the center of the geographical distribution of the parental population. Within the Caribbean population, we find that the proportion of Venezuela ancestry decreases slightly but not significantly as the distance from Caracas increases (ρ_Spearman_= −0.3562, p-value = 0.088; Figure 4). These results suggest that the Caribbean populations are akin to a contact zone between the Arizona and Venezuela populations. More generally, the results collectively show that in spite of the strong differentiation between *C. posadasii* populations there is evidence of admixture and gene exchange.

**FIGURE 4.**
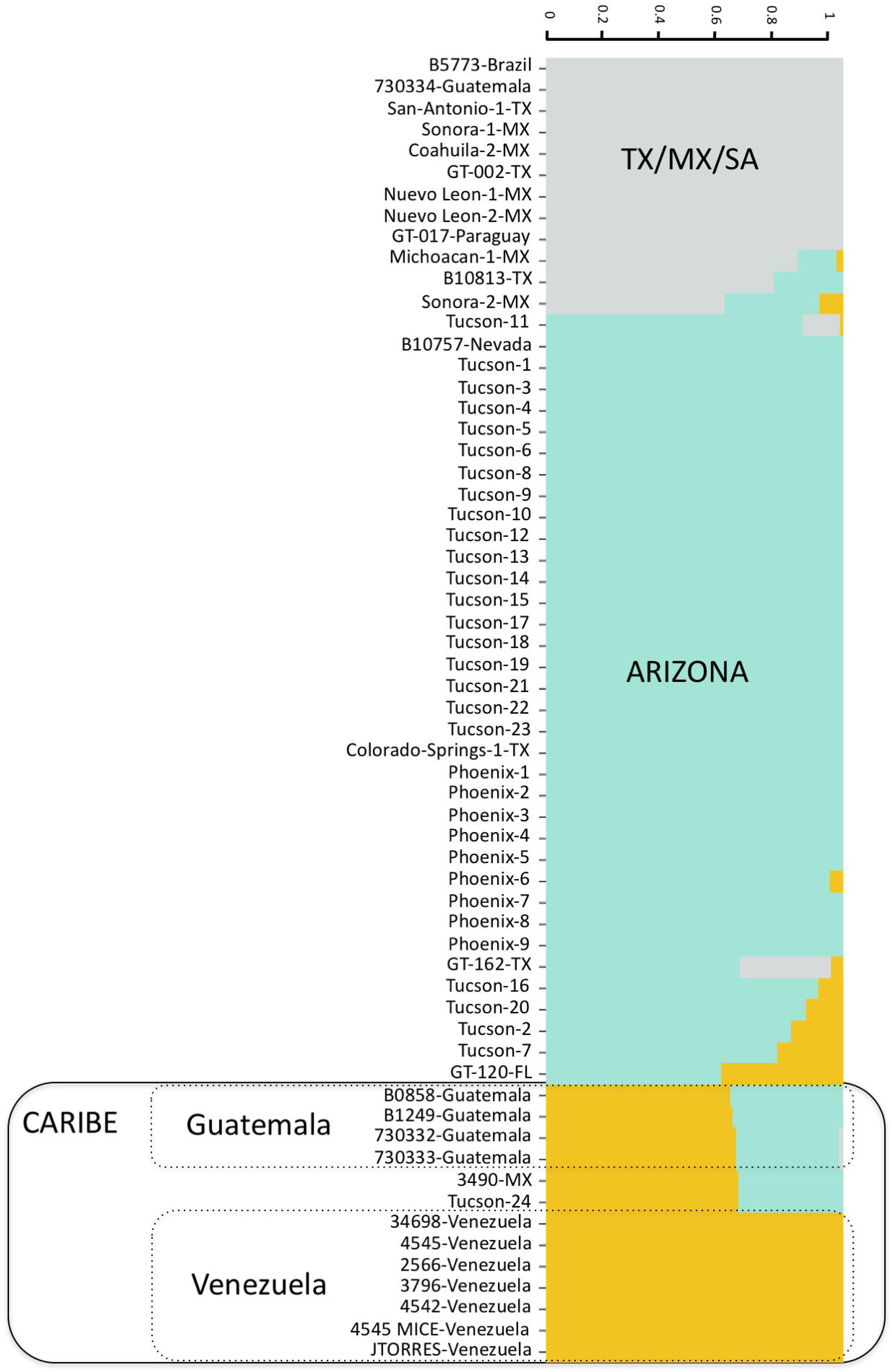
Within species admixture in *C. posadasii*. (A)Individual admixture proportions of *C. posadasii* isolates as inferred with ADMIXTURE. The height and colors of percentage of each population represent the probability of a given strain belonging to a given population. (B) The proportion of Venezuela admixture decreases as the distance from Caracas increases.

## DISCUSSION

In this study, we assess the magnitude of the differentiation within *C. posadasii* using whole genome sequence comparisons. The magnitude and partitioning of genetic variation in human fungal pathogens is a key, yet underappreciated, aspect of pathogen biology. *Coccidioides* represents a clear example of this paucity. Even though coccidioidomycosis exists in Central and South America, very little is known about the biology of the fungus in this region. Preliminary population assessments indicated that isolates from Central America belong to *C. posadasii* but these are genetically differentiated from both the *C. posadasii* TX/MX/SA clade and the Arizona clade (22). Notably, little is known in terms of the genetic diversity of isolates from xeric environments from South America, which poses the possibility that this snapshot of the relationships between *C. posadasii* isolates is incomplete.

Epidemiological studies show that coccidioidomycosis in South America is mostly caused by *C. posadasii* (19, 61, 64). Molecular analyses of soil DNA revealed that *Coccidioides* is common in xeric environments of Venezuela: all sampled sites (N = 15) were positive for *Coccidioides;* and sequencing of one of the ribosomal Internal Transcribed Spacers (ITS2) suggested that all the environmental samples from the region are, indeed, *C. posadasii* (33). Nonetheless, the sample of genotypes was diverse and encompassed multiple ITS2 haplotypes. If *C. posadasii* is so common in the soil in Venezuela, why is coccidioidomycosis so rare in the same area? One possibility is that the *C. posadasii* Venezuela lineage is less infective than other *Coccidioides* lineages. Since *C. posadasii* in Venezuela seems to have undergone a strong bottleneck (29), it is possible that the fungus has accumulated deleterious mutations that diminish its ability to establish infection in humans. Another possibility is that the Venezuela lineage is less virulent than other lineages. In this scenario, even if the fungus comes into contact with humans, and has the same ability to cause infection as other lineages, the disease is more likely to be asymptomatic than in other lineages. This would be consistent with the fact that large proportion of the people in Venezuela react to coccidioidin. The proportion of reactors in Venezuela is comparable to other human populations (65, 66). Variation in the ability to cause disease, either because of genotypic differences in the ability to infect or to induce disease, has been reported for other fungi of the Onygenales. *Histoplasma* spp., for example, show differences in fungal burden, disease kinetics, symptomology, and cytokine responses in controlled infections (67). Currently there is no definitive evidence of these differences among lineages of *Coccidioides*. It is also possible that coccidioidomycosis is underreported in Venezuelan (and other Latin American) populations compared to USA populations. Additionally, arid regions in South America tend to be associated with lower population densities and lower socio-economic status than coastal cities. Certainly, all these possibilities are not mutually exclusive. Defining the relative importance of these factors that may contribute to the low numbers of coccidioidomycosis in South America will required a combination of population genetics, genome-wide association studies, and fine-scale and comparable epidemiological studies paired with environmental surveys in multiple *Coccidioides* populations from different regions of endemicity.

One proposed hypothesis to explain *C. posadasii* genotypic distributions in South America is dispersion associated with human migration from North America (61). Early settlements of humans in Venezuela (Taima-Taima archeological site, Falcon state) were dated to 15k years ago (68). The divergence between the Venezuela and North America lineages of *C. posadasii* goes back to at least ~145,000-390,000 years ago, within the Middle and Upper Pleistocene (69). The confidence intervals of these dates do not overlap with the more well accepted understanding of human colonization of the Americas (less than 17,000 years ago; (70–72) but see (73)). Nonetheless, there are several caveats and alternative hypotheses that need to be taken into account. One possibility is that the mutation rate, which was used to convert genetic differentiation to years of divergence, differs across multiple groups of *Coccidioides*. The difference in mutation rates between *Coccidioides* lineages would have to high (~10X) to qualitatively change our conclusions. Second, it is possible the Venezuela lineage originated somewhere else (i.e., North America) and went extinct in its location of origin. If this lineage was only able to establish itself in Venezuela, then the hypothesis of human-driven migration of *C. posadasii* to South America would be compatible with our results. Third, other animal migrations and host-pathogen associations have not been explored. The Great American Biotic Interchange resulted in many species that could be preferred or novel host species moving from North America to South America and vice versa. Finally, the vast majority of isolates analyzed to date are derived from human patients and we cannot be fully confident of their site of origin or of overall genetic diversity in the environment. The patterns of dispersion of fungal pathogens, including *C. posadasii*, remain an underexplored topic.

The study of population structure in fungi has been almost exclusively applied to defining cryptic species. These lineages, which show the genetic signature of reproductive isolation (i.e., the phylogenetic species concept, reviewed in (74)), are morphologically similar and were not known to be distinct before DNA sequencing became possible (reviewed in (75)). This approach has led to the conclusion that the number of species of fungi is significantly underestimated as each species recognized by morphology harbors several phylogenetic species (4). Delineating species boundaries has important consequences for epidemiology and our understanding of pathogenesis; human pathogens previously assigned to a single species have been found to show different levels of virulence and in some cases cause different diseases. The case of *Histoplasma* exemplifies this utility. Recently, four species of *Histoplasma* were formally adopted following genomic differentiation. These lineages differ in their genome size, in their ability to cause disease and their geographical distribution (76). Analyses of gene flow show that these species exchange genes rarely but that most introgressions are found at low allele frequency which in turn might indicate the possibility that these exchanged alleles are deleterious, or slightly deleterious.

The magnitude of genetic differentiation seems to be insufficient to describe the different lineages of *C. posadasii* as well-formed different species. The case of the Venezuela clade in which the magnitude of interpopulation genetic distance is higher than the variation within Venezuela is worth highlighting. This pattern is caused by two main factors. First, the Venezuela lineage is differentiated from other populations. On average, the ratio of mean interpopulation distance to intrapopulation variation (outside Venezuela) is 1.2. Second, the Venezuela lineage shows a reduction in genetic diversity of 50X compared to other *C. posadasii* populations. These two patterns have different biological interpretations. First, there is strong population structure between species of *Coccidioides* which in turn might be a signal of incipient and recent speciation. Nonetheless, differentiation between incipient speciation and strong population structure is a major challenge (77, 78), which cannot be tackled without *prima facie* evidence of reproductive isolation. Additionally, the lower variability in Venezuela leads to a skewed ratio of inter-to intra population variation in pairwise comparisons involving Venezuela, and to some extent Guatemala, but this is not caused by an increase in the interpopulation differentiation but by an extreme reduction in intraspecific variation. Moreover, we observe a single mating-type *(MAT1-2)* within the Venezuelan population and according to the typical bipolar mating system of Eurotiomycetes, both opposite mating types cells *(MAT1-1* and *MAT1-2)* are need for sexual recombination. Thus, we conclude that this is likely an extreme bottleneck event, and not a signature of speciation.

Our analyses are parallel to other approaches previously used to quantify the magnitude of interspecific gene flow between species of pathogenic fungi, but our focus differs from these studies; our goal was to study whether populations of *C. posadasii* across the geographic range of the species are differentiated and exchange alleles. An overlooked aspect of population genetics in fungi is the magnitude of population structure within species. Understanding how species are separated into populations across geography is a prerequisite to address where genetic variation associated with virulence is originating across the geographic range of a species. Intraspecific population structure and speciation are part of a continuum that is modulated by the strength of reproductive isolation and the extent of genetic divergence between lineages (79, 80). The continuous nature of divergence and speciation makes the distinction of species boundaries challenging because population genetics estimates of differentiation cannot distinguish between nascent species and well-structured and old populations of the same species (78, 81). This is especially true for populations that occur in allopatry and do not interbreed.

Studying the magnitude of intraspecific gene exchange is not devoid of challenges. Species that have achieved genome-wide reciprocal monophyly will show fixed differences, which allows for detection of alleles that have crossed species boundaries after hybridization (e.g., (28, 82)). Populations from the same species, or even incipient species, are less likely to show these fixed differences, and analyses of introgression must be done based on differences of allele frequencies. This approach has proven useful in human populations (83) and maize (84) (reviewed in (85)). In the case of *C. posadasii* populations, it is possible that shared ancestry among populations is not caused by gene exchange but by incomplete lineage sorting (ILS) across different populations. This retention of ancestral alleles might be due to balancing selection, or simply by chance (86, 87). As genetic divergence accumulates the likelihood of retaining ancestral polymorphism by chance decreases, thus when divergence is still recent (populations or incipient species) ILS should be common (87, 88).

The *Coccidioides posadasii* population from Venezuela is not the only fungal pathogen from this country that shows signatures of genetic isolation. In the case of *Paracoccidioides*, two species coexist in the same geographic range; one endemic species, *P. venezuelensis*, and a species with broader distribution, *P. americana*, found in Venezuela and Brazil (89, 90). Whether the environments of Venezuela facilitate divergence, or whether it is a case of differences in sampling effort remains unknown. These two possibilities are both likely as xeric environments in South America have not been systematically sampled for *Coccidioides*, and it is possible that each xeric environment harbors its own lineage of *Coccidioides*. In the case of *Paracoccidioides*, sampling effort has been roughly equivalent in multiple countries and Venezuela harbors two species, a number similar to the number of species in Brazil, a country with an area nine times larger.

Hybridization and gene exchange seem to be of common occurrence during the evolutionary history of fungal pathogens. For example, *Paracoccidioides* species show complete mitochondrial capture which is reflected in the discordance between nuclear and mitochondrial gene genealogies (50). *Histoplasma ohiense* and *Histoplasma mississippiense* show evidence of admixture in their nuclear genome but such exchanged alleles are at low frequency (76, 82). *Cryptococcus* species also show evidence of shared genetic variation (91). This pattern is not limited to human pathogens. Plant pathogens and saprobic fungi also show evidence of gene exchange but the features that govern the magnitude of gene exchange remain unknown (e.g., (92, 93); reviewed in (94)). These general patterns are consistent with trends observed in other taxa but need to be formally tested in fungi.

Genetic variation among all taxa is dictated by the generation of new variants by mutation, migration, and recombination (63, 95). Geography plays a fundamental role on how these variants are maintained over time (96). Semi-isolated lineages might serve as reservoirs of variation which might feed into the main gene pool of a species by regular, but not constant, gene flow. These metapopulation dynamics in which populations are connected to each other, but where there is the possibility for population divergence and contact, is not exclusive to pathogens. In the case of *Coccidioides*, a systematic sampling of xeric and other suitable environments is sorely needed to assess how selection might dictate the sojourn of new mutations and also the evolutionary history of the species. More generally, approaches that incorporate spatial and temporal partitioning of genetic variation in pathogens will be crucial to understanding the factors that shape the genome and species history of organisms crucial to human well-being.

## Acknowledgments

We would like to thank Dr. Mireya Mendoza for initiating the Venezuelan *Coccidioides* project, and we wish her the best in her retirement. Technical sequencing support from TGen-North (Flagstaff, AZ) was provided by Dr. Dave Engelthaler. This work was supported by NIH/NIAID award R21AI28536 to BMB and NIH/NIGMS award R01GM121750 to DRM. The authors declare no conflicts of interest.

---

FIGURE S1. Cross-validation error learning curve of the Loglikelihood values collected by ADMIXTURE from k=1 to k=8 population scenarios.

TABLE S1. SRA accession number, isolate identifier, species and origin of isolation for each *Coccidioides* strain used for phylogenomics and population genetic analyses

TABLE S2. Genetic distances between all pairwise population comparisons within *C. posadasii*. We used asymptotic 2-sample Permutation Tests to compare intra- and interspecific differences.

